# Le Petit Prince (LPP) Multi-talker: Naturalistic 7T fMRI and EEG Dataset

**DOI:** 10.1101/2025.02.26.640265

**Authors:** Qixuan Wang, Qian Zhou, Zhengwu Ma, Nan Wang, Tianyu Zhang, Yaoyao Fu, Jixing Li

**Author notes:** (first). (co-first).

## Abstract

Prior neuroimaging datasets using naturalistic listening paradigms have predominantly focused on single-talker scenarios. While these studies have been invaluable for advancing our understanding of speech and language processing in the brain, they do not capture the complexities of real-world multi-talker environments. Here, we introduce the “Le Petit Prince (LPP) Multi-talker Dataset”, a high-quality, naturalistic neuroimaging dataset featuring 40 minutes of electroencephalogram (EEG) and 7T functional magnetic resonance imaging (fMRI) recordings from 26 native Mandarin Chinese speakers as they listened to both single-talker and multi-talker speech streams. Validation analyses conducted on both EEG and fMRI data demonstrate the dataset’s high quality and robustness. Additionally, the dataset includes detailed transcriptions and prosodic and linguistic annotations of the speech stimuli, enabling fine-grained analyses of neural responses to specific linguistic and acoustic features. The LPP Multi-talker Dataset is well-suited for addressing a wide range of research questions in cognitive neuroscience, including selective attention, auditory stream segregation, and working memory processes in naturalistic listening contexts.

## Background & Summary

Prior neuroimaging datasets employing naturalistic listening paradigms have predominantly focused on single-talker speech as stimuli^1–5^. This single-talker listening paradigm has been instrumental in identifying brain regions involved in various aspects of auditory and linguistic processing, including acoustic and phonemic analysis^6^, semantic comprehension^7^, and syntactic parsing^8^. However, everyday communication often unfolds in dynamic, multi-talker environments. In the well-known “cocktail party” scenarios^9^, listeners must selectively attend to a single speech stream, extract relevant information from background chatter, and adapt to rapidly shifting acoustic and linguistic cues^10^. These cognitive demands engage a broader range of neural processes beyond those typically activated in single-talker scenarios. Prior studies have demonstrated that listeners with normal hearing can selectively focus on a target speaker in the presence of competing speakers^11,12^. This selective attention enhances neural responses to the attended speech stream^13–18^.

Despite the significance of multi-talker scenarios for understanding speech processing in the brain, open neuroimaging datasets utilizing naturalistic multi-talker paradigms remain scarce. To address this gap, we introduce the “Le Petit Prince (LPP) Multi-talker Dataset”, a high-quality, multimodal neuroimaging dataset that captures neural responses to both single- and multi-talker speech streams. The LPP Multi-talker Dataset features both electroencephalogram (EEG) and 7T functional magnetic resonance imaging (fMRI) recordings from 26 native Mandarin Chinese speakers as they listened to both single-talker and multi-talker speech streams. EEG offers millisecond-level temporal resolution, enabling the study of rapid neural oscillations and event-related potentials associated with real-time speech processing. 7T fMRI provides a higher signal-to-noise ratio (SNR) and spatial resolution compared to 3T fMRI^19^, allowing for precise tracking of the neural dynamics of linguistic processing at fine-grained anatomical detail.

Validation analyses confirm the dataset’s high quality and robustness across participants, with clear differentiation between single-talker, attended and unattended speech in multi-talker conditions in bilateral temporal brain regions that are critical for speech and linguistic processing^20^. The dataset also includes comprehensive annotations for the auditory stimuli, covering speech transcriptions and detailed word-level and phrase-level annotations, such as log frequency, part-of-speech (POS) tags, and the number of parser actions for each word derived from Stanford parser^21^ based on bottom-up, top-down, and left-corner parsing strategies. These rich annotations enable fine-grained analysis of how specific acoustic and linguistic properties influence neural processing in complex auditory environments. All data and annotations are provided in standardized formats to ensure accessibility and reproducibility (see Fig. 1 for a detailed schematic overview of the data collection procedures, preprocessing steps, technical validation of the neuroimaging data, and annotation processes).

**Fig. 1.**
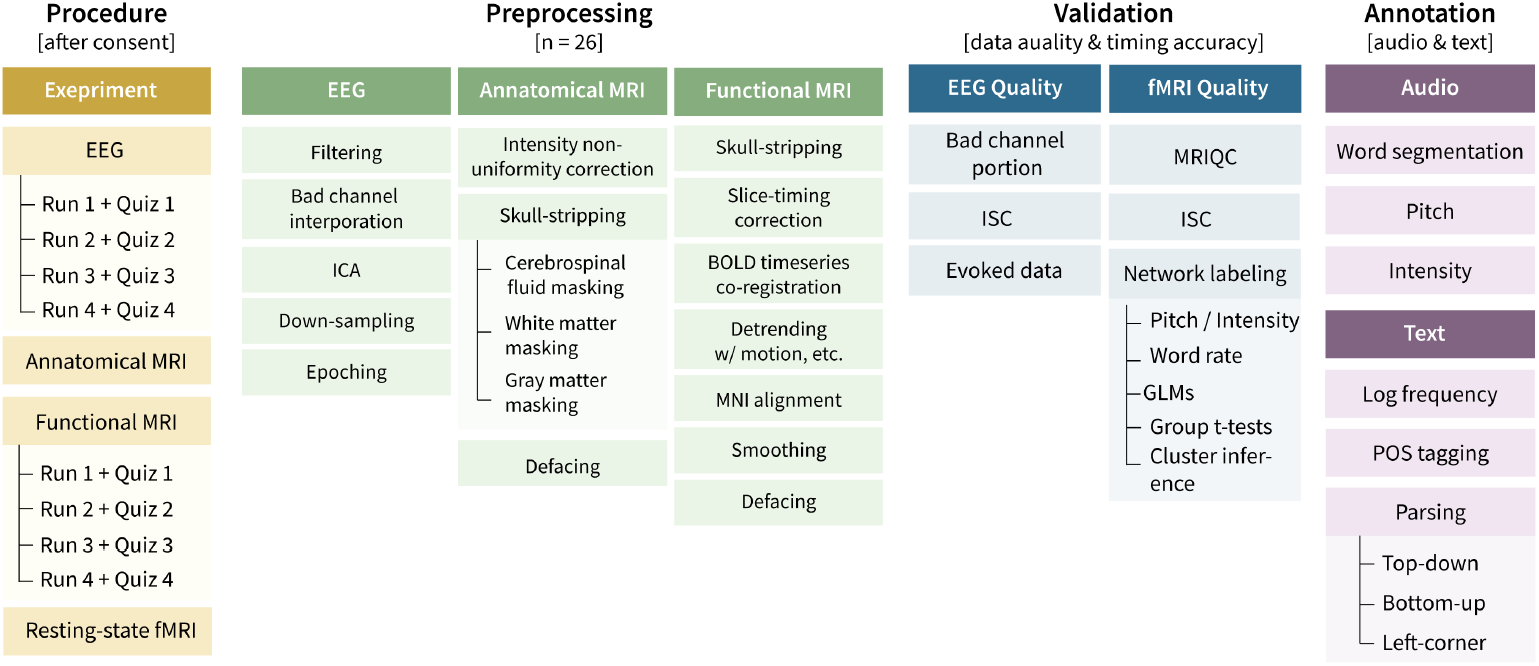
Schematic overview of the LPP-Multi-talker data collection procedures, preprocessing, technical validation and annotation. The data collection procedure (brown) involved recording neuroimaging signals while participants listened to four sections of an audiobook. EEG data were recorded approximately two years prior to the fMRI data using the same experimental design. The fMRI data collection process included anatomical MRI, followed by functional MRI, and resting-state fMRI. After data collection, preprocessing (green) was carried out, followed by behavioral and overall data quality assessments (blue). Audio and text annotations were generated using NLP tools (purple).

One of the key strengths of the LPP Multi-talker Dataset lies in its ecological validity. Unlike highly controlled laboratory paradigms, this dataset emulates real-world multi-talker listening conditions, providing a robust resource for investigating the neural mechanisms underlying selective attention, auditory stream segregation, and adaptive listening—processes essential for everyday communication. Additionally, the dataset supports cross-disciplinary research, offering a valuable resource for testing brain-computer interface (BCI) applications aimed at neural speech decoding in complex multi-talker environments.

## Methods Participants

A total of 26 participants (15 females, mean age=23.96±2.23 years) took part in both the EEG and fMRI studies. All participants were right-handed native Mandarin speakers enrolled in an undergraduate or graduate program in Shanghai and had no self-reported history of neurological disorders. EEG and fMRI recordings were collected while the participants listened to two sections of the Chinese version of “Le Petit Prince”, narrated by one or two speakers simultaneously. The fMRI data were collected two years after the EEG data collection to reduce the potential interference of prior exposure to the experimental material (mean interval=957±45 days; see Table 1 for the participants’ demographic information and their data acquisition time). All participants were provided with written informed consent prior to the experiments and were paid for their participation.

**Table 1.** Participants’ demographic information and their EEG and fMRI data acquisition time.

| Participant ID | Sex | Age | EEG Acquisition Time | fMRI Acquisition Time |
| --- | --- | --- | --- | --- |
| sub-01 | F | 26 | 2021-09-10 | 2024-07-05 |
| sub-02 | F | 23 | 2021-09-15 | 2024-06-24 |
| sub-03 | M | 24 | 2021-09-16 | 2024-06-27 |
| sub-04 | F | 21 | 2021-09-18 | 2024-06-14 |
| sub-05 | F | 21 | 2021-09-28 | 2024-06-28 |
| sub-06 | F | 24 | 2021-10-31 | 2024-05-30 |
| sub-07 | M | 27 | 2021-11-07 | 2024-06-21 |
| sub-08 | M | 20 | 2021-11-07 | 2024-05-28 |
| sub-09 | M | 25 | 2021-11-10 | 2024-06-25 |
| sub-10 | M | 23 | 2021-11-13 | 2024-06-14 |
| sub-11 | M | 28 | 2021-11-14 | 2024-06-18 |
| sub-12 | F | 26 | 2021-12-18 | 2024-05-31 |
| sub-13 | F | 24 | 2021-12-14 | 2024-04-09 |
| sub-14 | F | 23 | 2021-12-13 | 2024-06-05 |
| sub-15 | F | 26 | 2021-09-12 | 2024-05-28 |
| sub-16 | F | 21 | 2021-09-19 | 2024-06-07 |
| sub-17 | F | 26 | 2021-09-22 | 2024-07-05 |
| sub-18 | F | 26 | 2021-09-09 | 2024-04-03 |
| sub-19 | M | 23 | 2021-09-26 | 2024-06-18 |
| sub-20 | F | 21 | 2021-10-31 | 2024-07-05 |
| sub-21 | M | 27 | 2021-11-04 | 2024-05-31 |
| sub-22 | M | 25 | 2021-09-10 | 2024-04-09 |
| sub-23 | M | 21 | 2021-12-18 | 2024-06-18 |
| sub-24 | M | 26 | 2021-12-18 | 2024-07-01 |
| sub-25 | F | 24 | 2021-12-16 | 2024-06-05 |
| sub-26 | F | 22 | 2021-11-20 | 2024-08-26 |

### Stimuli

The stimuli were the same for both EEG and fMRI experiments and consist of two sections of a Chinese translation of the novel “The Little Prince” (available at http://www.xiaowangzi.org/). The two excerpts were narrated by one male and one female computer-synthesized voice, developed by the Institute of Automation, Chinese Academy of Sciences. The synthesized speech (available at https://osf.io/fjv5n/) is comparable to human narration, as confirmed by participants’ post-experiment assessment of its naturalness. Additionally, using computer-synthesized voice instead of human-narrated speech alleviates the potential issue of imbalanced voice intensity and speaking rate that can arise between female and male narrators. The two sections were matched in length (approximately 10 minutes) and mean amplitude (approximately 65 dB) and were mixed digitally in a single channel to prevent any biases in hearing ability between the left and right ears.

### Experiment procedures

The experimental task consists of a multi-talker condition and a single-talker condition (see Fig. 2a). In the multi-talker condition, the mixed speech will be presented twice, with the female and male speakers narrating simultaneously. Before each trial, instructions appeared in the center of the screen indicating which of the talkers to attend to (e.g., “Attend Female”). In the single-talker condition, the male and female speeches were presented separately. The presentation order of the four conditions were randomized. For the EEG experiment, participants rated the intelligibility of the multi-talker and single-talker speeches on a 5-point Likert scale at the end of the experiment. For the fMRI experiment, participants completed four quiz questions after each run through a button box (see Table S1 in Supplementary for all quiz questions). These questions were used to confirm their comprehension and will be viewed by the participants via a mirror attached to the head coil. Stimuli were presented using insert earphones (ER-3C, Etymotic Research, United States) for the EEG experiment and MRI-compatible headphones (OptoACTIVE™ Active Noise Control Optical MRI Communication System, Optoacoustics Ltd., Israel) for the fMRI experiment. Participants were instructed to maintain visual fixation for the duration of each trial on a crosshair centered on the screen, and to minimize eye blinking and all other motor activities for the duration of each section (see Table S2 in Supplementary for the run order of the participants). The whole experiment, including preparation time and practice, lasted for around 60 minutes. The experimental procedures have been approved by the Ethics Committee of City University of Hong Kong and the Ninth People’s Hospital affiliated with Shanghai Jiao Tong University School of Medicine (EEG: SH9H-2019-T33-2; fMRI: SH9H-2022-T379-2).

**Fig. 2.**
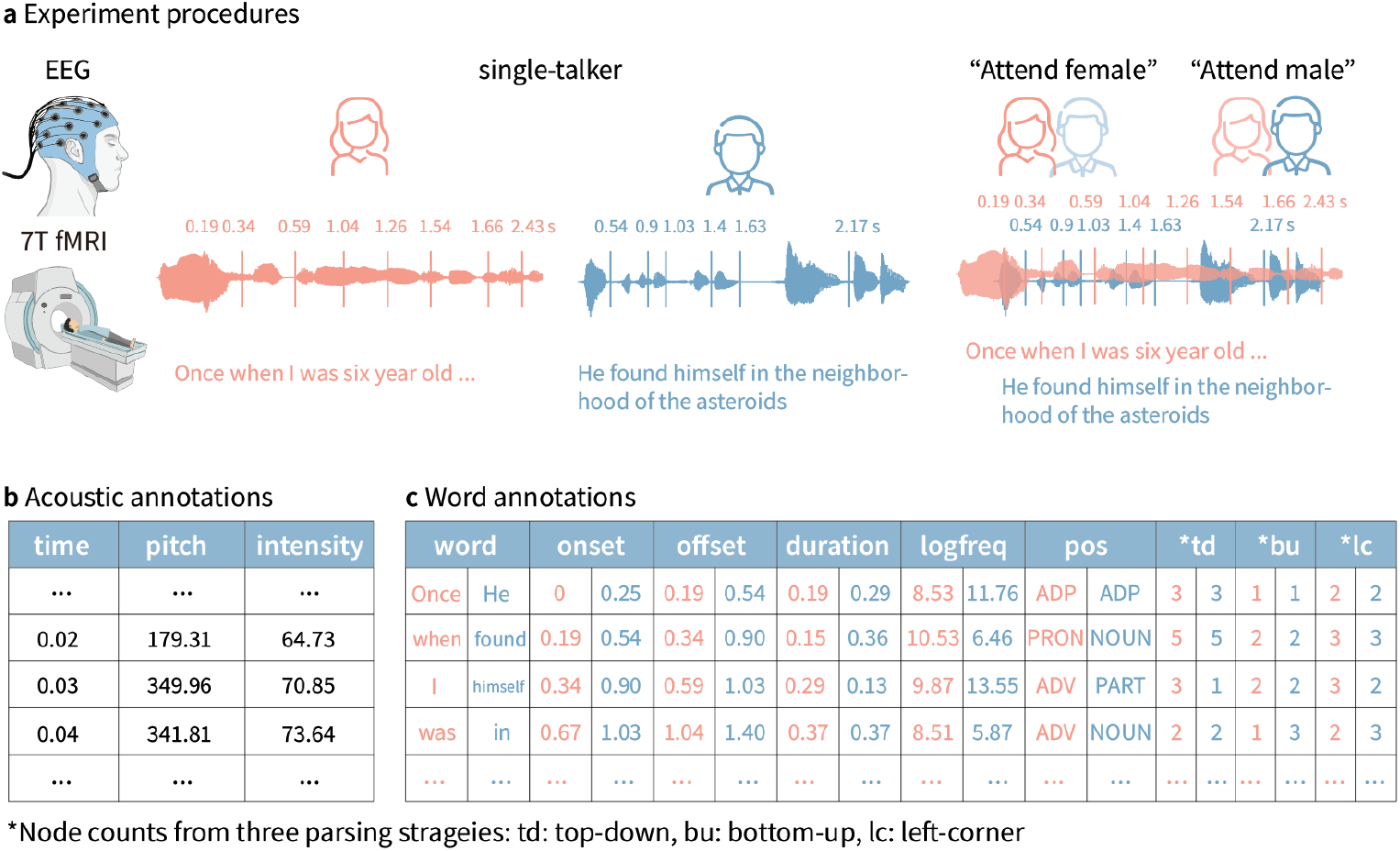
Experiment procedures and annotations. (a) The experiment consisted of two multi-talker conditions and two single-talker conditions. In the multi-talker condition, speech was presented by a female and a male speaker, each narrating a different section of the audiobook simultaneously. Prior to each section, instructions appeared at the center of the screen indicating which speaker participants should attend to (e.g., “Attend female”). In the single-talker condition, the male and female speeches were presented separately. The stimuli were delivered to participants in a random order. (b) The fundamental frequency (pitch) and intensity of the audio stimuli were calculated. (c) Word segmentation, part-of-speech tagging, log-transformed word frequency, and node counts were obtained using top-down, bottom-up, and left-corner parsing strategies.

### Data acquisition

The EEG data was collected at the Department of Otolaryngology-Head and Neck Surgery, Shanghai Ninth People’s Hospital affiliated with the School of Medicine at Shanghai Jiao Tong University. EEG was recorded using a standard 64-channel actiCAP mounted according to the international 10-20 system against a nose reference (Brain Vision Recorder, Brain Products). The ground electrode was set at the forehead. Signals were recorded with a sampling rate of 500 Hz, and the frequency range was set between 0.016 Hz and 80 Hz. Impedance levels were maintained below 20 kΩ.

The fMRI data was collected in a 7.0 T Terra Siemens MRI scanner at the Zhangjiang International Brain Imaging Centre, located at the Institute of Science and Technology for Brain-Inspired Intelligence, Fudan University, Shanghai. Anatomical scans were obtained using a Magnetization Prepared RApid Gradient-Echo (MP-RAGE) SAG iPAT2 pulse sequence with T1-weighted contrast (256 single-shot interleaved sagittal slices with A/P phase encoding direction; voxel size=0.7×0.7×0.7 mm; FOV=208 mm; TR=3800 ms; TE=2.32 ms; flip angle=7°; acquisition time=3 s; GRAPPA in-plane acceleration factor=3). Functional scans were acquired using T2-weighted echo planar imaging (85 interleaved axial slices with A/P phase encoding direction; voxel size=1.6×1.6×1.6 mm; FOV=208 mm; TR=1000 ms; TE=22.2 ms; multiband acceleration factor for parallel slice acquisition=5; flip angle=45°).

### Data preprocessing

We applied a minimal preprocessing pipeline on the EEG data using MNE-Python (v1.4.2; Gramfort et al., 2013). We first identified and interpolated bad channels using the PyPREP package (v0.4.3). We then band-pass filtered the EEG data between 0.1 Hz and 39 Hz using a linear-phase finite impulse response (FIR) filter. We applied independent component analysis (ICA) to remove eye blink artifacts using the FastICA algorithm. The data were subsequently downsampled to 100 Hz and were segmented into four epochs corresponding to the four conditions, with each epoch spanning from -500 ms to 10 minutes after each section onset.

For the fMRI data, all digital imaging and communications in medicine (DICOM) images were converted to the brain imaging data structure (BIDS) using dcm2bids (v3.1.1^22^) and then converted to neuroimaging informatics technology initiative (NIfTI) format using dcm2niix (v1.0.20220505). Anonymization of participants was then performed with PyDeface (v2.0.2), which removes voxels associated with facial features. Preprocessing of the neuroimaging data was conducted using fMRIPrep (v20.2.0^23^. For the anatomical data, T1-weighted images were corrected for intensity non-uniformity, skull stripping, and segmentation into gray matter (GM), white matter (WM), and cerebrospinal fluid (CSF). These images were then registered to Montreal Neurological Institute (MNI) space using the MNI152NLin2009cAsym:res-2 template, ensuring consistent spatial normalization across participants. Functional MRI preprocessing included skull stripping, motion correction, slice-timing correction, and co-registration to the T1-weighted reference. The blood-oxygen-level-dependent (BOLD) time series were resampled to both native and MNI space, and various confound time series were calculated. Noise correction was applied to improve the signal-to-noise ratio, and motion outliers were identified to account for residual artifacts in subsequent analyses.

### Annotations

Annotation of the speech stimuli includes prosodic information, time-aligned word segmentation and linguistic predictors, from lexical to syntactic levels, using freely-available natural language processing (NLP) tools. Fundamental frequency (f0) and intensity of the audio was extracted at 10 ms intervals using Praat-Parselmouth (v0.4.4^24^). Log-transformed unigram frequency of each word in the stimuli was estimated using the Google Books Ngram Viewer dataset. Part-of-speech (POS) tags for each word in the stimuli were assigned using the Stanford Parser^21^. Additionally, the number of parsing actions required for processing each word within its sentence context were derived from the Stanford constituency trees, based on bottom-up, top-down, and left-corner parsing strategies (see Fig. 2b for the annotation of an example sentence).

### Data Records

We anonymized all data by eliminating any information and anatomical details that could be used to identify the participants. The resulting files can be accessed from the OpenNeuro repository (https://openneuro.org/ds005345). Please refer to Fig. 3 for an overview of folder structure. A full description of the available content is included in the README file within the repository.

**Fig. 3.**
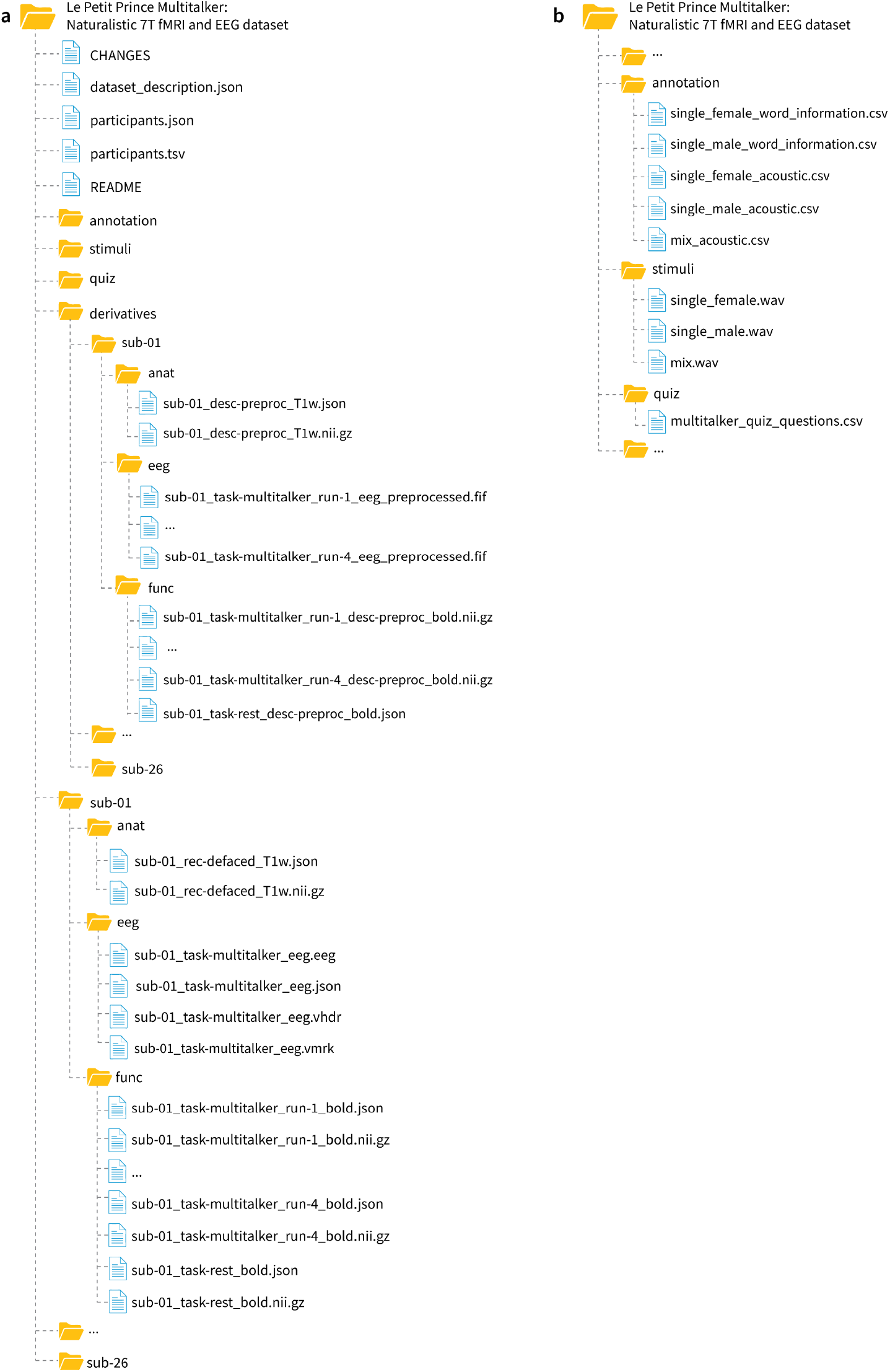
Overview of the folder structure. (a) Directories for the EEG and fMRI data. (b) Directories for the annotation of acoustic and word information, experimental stimuli and the quiz questions.

### Annotation files

Location: annotation/single_female[single_male, mix]_acoustic.csv annotation/single_female[single_male]_word_information.csv

File format: comma-separated value.

Annotation of acoustic and linguistic features for the audio and text of the stimuli.

### Audio files

Location: stimuli/single_female[single_male, mix].wav

File format: wav.

The 10-minute audio from the Chinese version of “Le Petit Prince”.

### Quiz file

Location: quiz/multi-talker_quiz_questions.csv File format: comma-separated value.

The 16 multiple choice questions employed in both the fMRI and EEG experiments.

### EEG data files

Location: sub-<ID>/eeg/sub-<ID>_task-multi-talker_eeg.eeg

File format:eeg, BrainVision Core Data Format

The preprocessed data are also available as:

derivatives/sub-<ID>/eeg/sub-<ID>_task-multi-talker_run-1[2-4]_eeg_preprocessed.fif

### Anatomical MRI files

Location: sub-<ID>/anat/sub-<ID>_rec-defaced_T1w.nii.gz

File format: NIfTI, gzip-compressed.

Sequence protocol: sub-<ID>/anat/sub-<ID>_rec-defaced_T1w.json

The preprocessed data are also available as:

derivatives/sub-<ID>/anat/sub-<ID>_desc-preproc_T1w.nii.gz

### Functional MRI files

Location: sub-<ID>/func/sub-<ID>_task-multi-talker_run-1[2–4]_bold.nii.gz

File format: NIfTI, gzip-compressed.

Sequence protocol: sub-<ID>/func/sub-<ID>_task-multi-talker_run-1[2–4]_bold.json

The preprocessed data are also available as:

derivatives/sub-<ID>/func/sub-<ID>_task-multi-talker_run-1[2–4]_desc-preproc_bold.nii.gz

### Resting-state MRI files

Location: sub-<ID>/func/sub-<ID>_task-rest_bold.nii.gz

File format: NIfTI, gzip-compressed.

Sequence protocol: sub-<ID>/func/sub-<ID>_task-rest_bold.json

The preprocessed data are also available as:

derivatives/sub-<ID>/func/sub-<ID>_task-rest_desc-preproc_bold.nii.gz

### Technical validation EEG data quality

Quality metrics for the EEG data included the proportion of bad channels and the number of ICA components removed to correct for artifacts such as eye movement. We also evaluated the consistency of neural responses across participants by calculating the mean inter-subject correlation (ISC) across all EEG sensors for all sections, which provides a measure of shared neural activity in response to the experimental stimuli.

### Portion of bad channels and excluded ICA components

We used the PyPREP package (v0.4.3) to identify EEG channels with poor signal quality ^25^ based on abnormal correlations with neighboring channels. Fig. 4a illustrated the sensor locations. The number of bad channels for each participant was summarized in Fig. 4b (left). 13 out of 26 participants had no bad channels, reflecting overall good signal quality. ICA Components identified as artifacts were manually inspected and removed to ensure the preservation of neural signals. The number of ICA components removed during preprocessing was calculated for each participant and is displayed in Fig. 4b (right).

**Fig. 4.**
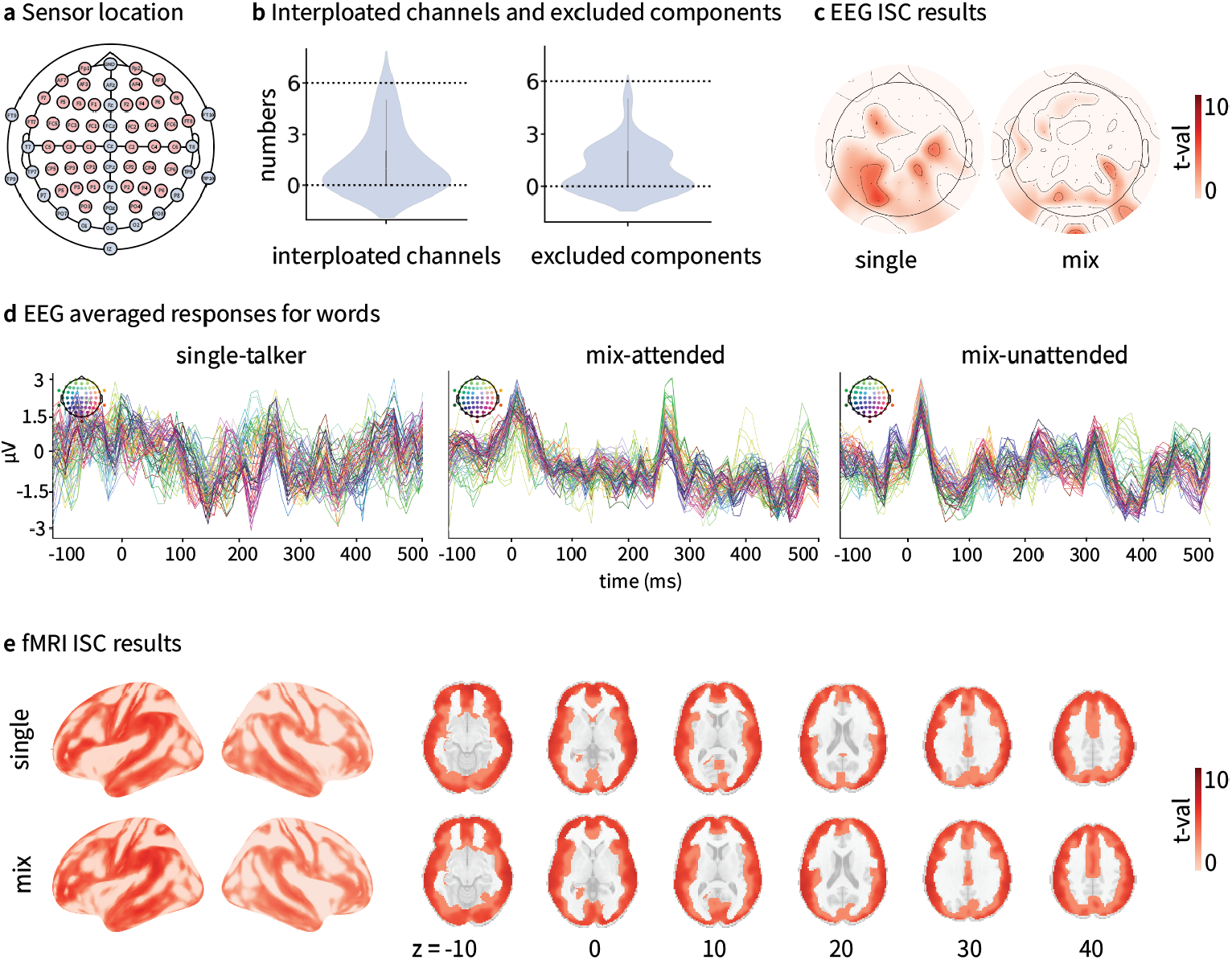
Assessment of EEG and fMRI data quality. (a) Sensor layout of the 64-channel EEG system. (b) Number of interpolated channels and excluded ICA components for each participant. (c) ISC results for EEG responses for the single-talker, attended, and unattended speech streams. (d) Evoked EEG responses for words in the single-talker, attendeded, and unattended speech streams. (e) ISC results for fMRI data under the single-talker and mixed-talker conditions.

### Inter-subject correlation (ISC) of EEG data

To evaluate the consistency of EEG responses across participants, we conducted an ISC^26^ analysis on the EEG data for all participants, for the single-talker and mixed-talker conditions separately. This analysis involved correlating the EEG time series for each participant with the average time series across all participants at each electrode and time point. Statistical significance of the ISC results at the group level was determined using a cluster-based one-sample, one-tailed t-test^27^. The analysis identified clusters of electrodes and time points where ISC values significantly exceeded chance levels (see Fig. 4c).

### Evoked responses for words

We segmented the EEG recordings into intervals spanning 100 ms before to 500 ms after each word offset, leading to 3,258 epochs per participant. The evoked responses for were computed across all participants and all 64 EEG channels over the 600 ms epoch, resulting in evoked responses for the single-talker, attended, and unattended speech streams (Fig. 4d). The analyses were performed with MNE (v1.4.2^28^).

### fMRI data quality

The fMRI data quality was evaluated using the MRI Quality Control tool (MRIQC). This included metrics assessing signal clarity, noise levels, and anatomical alignment. We also computed ISC of the fMRI data to evaluate the consistency of brain activity patterns across participants during the experiments. To further analyze the neural responses, we conducted whole-brain general linear modeling (GLM) using pitch, intensity, and word rate as regressors. These analyses were performed separately for the single-talker, attended, and unattended conditions for the dual-talker conditions. The results of these analyses aligned with findings from prior studies^2,3,15^.

### ISC of fMRI data

To evaluate the consistency of brain signal responses to stimuli under single-talker and mixed-talker conditions, we computed the ISC for each voxel’s time series across participants, following the methodology outlined by Li et al. (2022). For each participant, the activity of each voxel was correlated with the average time series of the corresponding voxel across all participants. This process was repeated for all participants, generating an ISC map at the group level. The ISC results demonstrated the highest correlations in the temporal regions for both single-talker and mixed-talker conditions (see Fig. 4e).

### MRIQC

We applied MRIQC to assess the quality of the fMRI data. The evaluation reports generated from image-quality metrics (IQMs) for the anatomical MRI (Fig. 5a) and functional MRI data (Fig. 5b) suggested overall high data quality: Low values of the coefficient of joint variation (CJV) for gray and white matter, along with the entropy-focus criterion (EFC), suggest minimal head motion and few artifacts. High contrast-to-noise ratio (CNR) and foreground-background energy ratio (FBER) indicate clear differentiation of tissue types and a well-balanced distribution of image energy. The white matter to maximum intensity ratio (WM2MAX) shows that the intensity of white matter is within the expected range. Additionally, the signal-to-noise ratio (SNR) for both gray and white matter indicates good overall data quality.

**Fig. 5.**
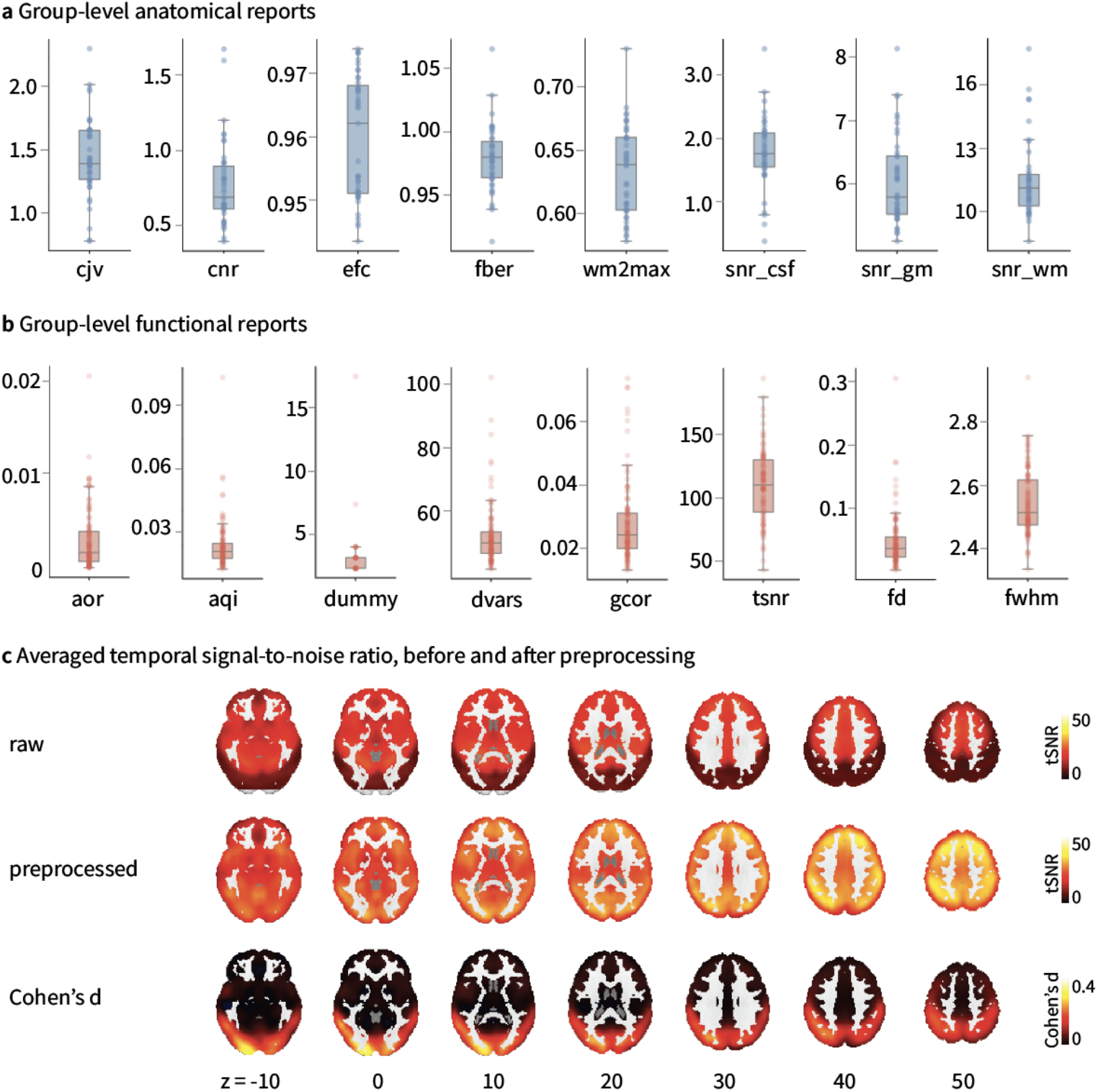
Assessment reports from the M. (a) Group-level IQMs of anatomical data. (b) Group-level IQMs of functional data. (c) Voxel-wise temporal tSNR analysis before and after preprocessing. Cohen’s *d* effect sizes indicated an increase in tSNR following preprocessing.

The functional MRI data also showed high temporal signal-to-noise ratio (tSNR) and low temporal derivative of time courses of root mean square over voxels (DVARS), suggesting strong temporal stability. Minimal head motion and artifacts are demonstrated by low average framewise displacement (FD) and AFNI’s outlier ratio (AOR). Moreover, low values of mean full width at half maximum (FWHM) and AFNI’s quality index (AQI) indicate a well-distributed and consistent image intensity across the brain. Low global correction (GOR) values and a small number of dummy scans suggest that the correction procedures were effective and the initial scan state was stable. We also compared the tSNR values for the raw and preprocessed functional MRI using Cohen’s d:

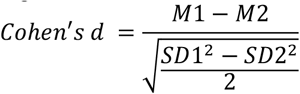

where *M* and SD represented the mean and standard deviation of tSNR within a voxel. A grey matter mask was applied to exclude white matter and subcortical areas. The results of Cohen’s d analysis revealed a substantial increase in tSNR following preprocessing, particularly within the occipital lobe (see Fig. 5c).

### General linear modeling (GLM) of acoustic features and word rate

Three GLMs were constructed to analyze fMRI data and assess the brain’s responses to pitch, intensity, and word rate across three experimental conditions: single-talker, attended, and unattended speech streams. Pitch and intensity at every 10 ms of the audio were extracted and convolved with the canonical hemodynamic response function (HRF) to create the acoustic regressors. The offset of each word in the audio was marked as one and convolved with the canonical HRF to create the word rate regressor. The time course of each voxel’s BOLD signal for each subject was modeled using the three regressors, and the resulting beta maps were tested for significance using one-sample t-tests at the group level. The threshold for cluster-level inference was set at p < 0.001 (see Fig. 6a for the procedure of the GLM analysis). The results showed significant bilateral temporal activity for the auditory and word rate regressors under the single-talker and attended speech condition, consistent with prior results (Li et al., 2022; Momenian et al., 2024) and the clusters obtained from meta-analyses of pitch, acoustic, and word using Neurosynth^29^. The unattended speech showed smaller cluster in the temporal regions, suggesting different neural responses to attended and unattended speech^2,3^ (see Fig. 6d for the significant brain clusters for pitch, intensity and word rate across the three listening conditions and Table S3 in Supplementary for detailed statistics of the clusters).

**Fig. 6.**
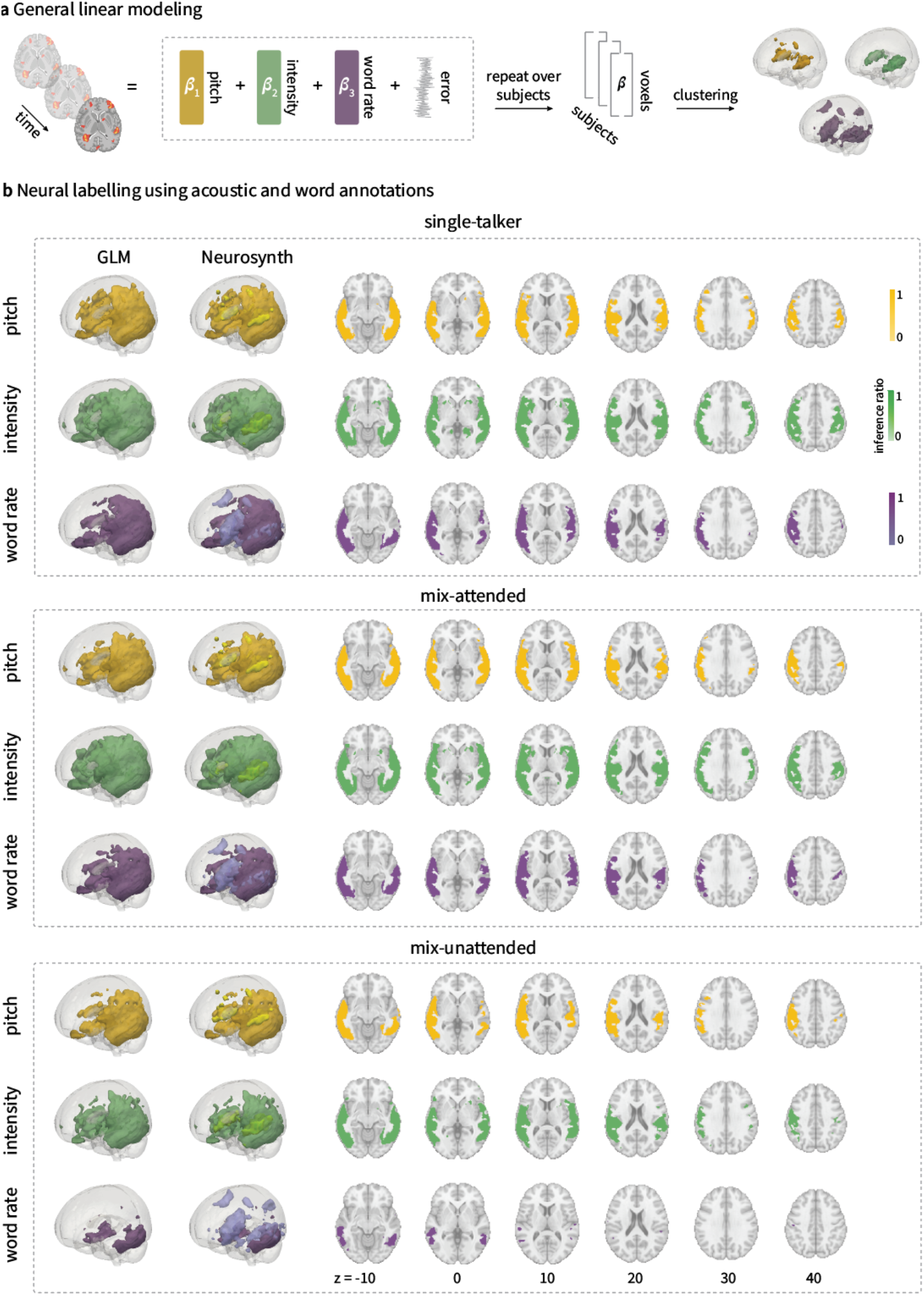
GLM analyses to localize pitch, intensity and word rate regressors of the single-talker, attended and unattended speech streams. (a) Overview of the GLM method. Pitch and intensity at every 10 ms of the audio were extracted and convolved with the canonical HRF to create the acoustic regressors. The offset of each word in the audio was marked as one and convolved with the canonical HRF to create the word rate regressor. The time course of each voxel’s BOLD signal for each subject was modeled using the three regressors, and the resulting beta maps were tested for significance using one-sample t-tests at the group level. The threshold for cluster-level inference was set at p < 0.001. (b) Significant clusters for the pitch, intensity and word rate under the single-talker, attended and unattended speech conditions, compared to the clusters obtained from meta-analyses of pitch, acoustic, and word using Neurosynth.

### Usage Notes

The multi-talker dataset we have developed offers valuable insights into speech comprehension in challenging conditions, such as the cocktail party scenario. However, there are several limitations and potential bottlenecks in its usage.

### Annotation Bottleneck

Although we made a clear distinction between “attended” and “unattended” conditions in the mixed-talker sessions, there is a possibility that participants might have adopted different attentional strategies oriented at “getting the gist” of the story during the mixed-speech condition or potentially disengage from the task due to comprehension difficulties. Although we have conducted to an ISC analyses on both the EEG and fMRI data to ensure consistency of neural responses across participants, it may still be the case that the annotations may not fully capture the cognitive and perceptual dynamics of the participants.

### Analysis bottleneck

Individual differences in cognitive abilities may lead to variations in how competing speech streams are processed, thereby introducing noise in group-level analyses. For example, a participant with higher working memory capacity might be able to maintain focus on the attended speaker despite the presence of distractors, whereas a participant with lower capacity may struggle to filter out irrelevant speech, resulting in neural patterns that do not align with the group as a whole.

In addition to univariate GLM analyses, more advanced analysis techniques, such as multivariate approaches or computational models that can jointly represent attended and unattended speech^15^, may be employed to further reveal the underlying neural mechanisms of selective attention and auditory processing, providing a more nuanced understanding of how the brain processes overlapping speech signals in real-world scenarios.

## Code availability

The dataset is publicly available at the OpenNeuro repository (https://openneuro.org/datasets/ds005345). The scripts can be accessed through GitHub (https://github.com/compneurolinglab/lpp_multi-talker).

## Acknowledgements

This work was supported by the National Natural Science Foundation of China (Wang Q. 82201273), the CityU Start-up Grant 7020086 and CityU Strategic Research Grant 7200747 (Li, J.).

## Author contributions

J.L. designed the study. Q.W. and Q.Z. collected the data. Z.M., N.W. and J.L. analysed data and wrote the paper. T.Z. and Y.F. helped write the paper.

## References

1. LeBel, A. et al. A natural language fMRI dataset for voxelwise encoding models. Sci Data 10, 555 (2023).

2. Li, J. et al. Le Petit Prince multilingual naturalistic fMRI corpus. Sci Data 9, 530 (2022).

3. Momenian, M. et al. Le Petit Prince Hong Kong (LPPHK): Naturalistic fMRI and EEG data from older Cantonese speakers. Sci Data 11, 992 (2024).

4. Nastase, S. A. et al. The “Narratives” fMRI dataset for evaluating models of naturalistic language comprehension. Sci Data 8, 250 (2021).

5. Wang, S., Zhang, X., Zhang, J. & Zong, C. A synchronized multimodal neuroimaging dataset for studying brain language processing. Sci Data 9, 590 (2022).

6. Gwilliams, L., King, J.-R., Marantz, A. & Poeppel, D. Neural dynamics of phoneme sequences reveal position-invariant code for content and order. Nat Commun 13, 6606 (2022).

7. Li, J., Lai, M. & Pylkkänen, L. Semantic composition in experimental and naturalistic paradigms. Imaging Neuroscience 2, 1–17 (2024).

8. Brennan, J. R., Stabler, E. P., Van Wagenen, S. E.Luh, W.-M. & Hale, J. T. Abstract linguistic structure correlates with temporal activity during naturalistic comprehension. Brain Lang 157–158, 81–94 (2016).

9. Cherry, E. C. Some Experiments on the Recognition of Speech, with One and with Two Ears. The Journal of the Acoustical Society of America 25, 975–979 (1953).

10. McDermott, J. H. The cocktail party problem. Curr Biol 19, R1024–1027 (2009).

11. Brungart, D. S. Informational and energetic masking effects in the perception of two simultaneous talkers. J Acoust Soc Am 109, 1101–1109 (2001).

12. Shinn-Cunningham, B. G. Object-based auditory and visual attention. Trends Cogn Sci 12, 182–186 (2008).

13. Brodbeck, C., Hong, L. E. & Simon, J. Z. Rapid Transformation from Auditory to Linguistic Representations of Continuous Speech. Curr Biol 28, 3976-3983.e5 (2018).

14. Ding, N. & Simon, J. Z. Emergence of neural encoding of auditory objects while listening to competing speakers. Proceedings of the National Academy of Sciences 109, 11854– 11859 (2012).

15. Li, J. et al. Multi-talker speech comprehension at different temporal scales in listeners with normal and impaired hearing. eLife 13, (2024).

16. Mesgarani, N. & Chang, E. F. Selective cortical representation of attended speaker in multi-talker speech perception. Nature 485, 233–236 (2012).

17. O’Sullivan, K., Dankaerts, W., O’Sullivan, L. & O’Sullivan, P. B. Cognitive Functional Therapy for Disabling Nonspecific Chronic Low Back Pain: Multiple Case-Cohort Study. Phys Ther 95, 1478–1488 (2015).

18. Zion Golumbic, E. M. et al. Mechanisms Underlying Selective Neuronal Tracking of Attended Speech at a ‘Cocktail Party’. Neuron 77, 980–991 (2013).

19. Ahveninen, J. et al. Intracortical depth analyses of frequency-sensitive regions of human auditory cortex using 7T fMRI. NeuroImage 143, 116–127 (2016).

20. Malik-Moraleda, S. et al. An investigation across 45 languages and 12 language families reveals a universal language network. Nat Neurosci 25, 1014–1019 (2022).

21. Levy, R. & Manning, C. D. Is it Harder to Parse Chinese, or the Chinese Treebank? in Proceedings of the 41st Annual Meeting of the Association for Computational Linguistics 439–446 (Association for Computational Linguistics, Sapporo, Japan, 2003). doi:10.3115/1075096.1075152.

22. Boré, A., Guay, S., Bedetti, C., Meisler, S. & GuenTher, N. Dcm2Bids. Zenodo 10.5281/zenodo.8342572 (2023).

23. Esteban, O. et al. fMRIPrep: a robust preprocessing pipeline for functional MRI. Nat Methods 16, 111–116 (2019).

24. Jadoul, Y., Thompson, B. & De Boer, B. Introducing Parselmouth: A Python interface to Praat. Journal of Phonetics 71, 1–15 (2018).

25. Bigdely-Shamlo, N., Mullen, T., Kothe, C.Su, K.-M. & Robbins, K. A. The PREP pipeline: standardized preprocessing for large-scale EEG analysis. Front. Neuroinform. 9, (2015).

26. Hasson, U., Nir, Y., Levy, I., Fuhrmann, G. & Malach, R. Intersubject synchronization of cortical activity during natural vision. Science 303, 1634–1640 (2004).

27. Maris, E. & Oostenveld, R. Nonparametric statistical testing of EEG- and MEG-data. Journal of Neuroscience Methods 164, 177–190 (2007).

28. Gramfort, A. et al. MEG and EEG data analysis with MNE-Python. Front. Neurosci. 7, (2013).

29. Yarkoni, T., Poldrack, R. A., Nichols, T. E., Van Essen, D. C. & Wager, T. D. Large-scale automated synthesis of human functional neuroimaging data. Nat Methods 8, 665–670 (2011).

